# Heritability of ethanol consumption and pharmacokinetics in a genetically diverse panel of Collaborative Cross mouse strains and their inbred founders

**DOI:** 10.1101/2020.09.13.294769

**Authors:** Jared R. Bagley, Elissa J. Chesler, Vivek M. Philip, Center for the Systems Genetics of Addiction, James D. Jentsch

## Abstract

**Background:** Inter-individual variation in voluntary ethanol consumption and ethanol response is partially influenced by genetic variation. Discovery of the genes and allelic variants that affect these phenotypes may clarify the etiology and pathophysiology of problematic alcohol use, including alcohol use disorder. Genetically diverse mouse populations also demonstrate heritable variation in ethanol consumption and can be utilized to discover the genes and gene networks that influence this trait. The Collaborative Cross (CC) recombinant inbred strains, Diversity Outbred (DO) population and their eight founder strains are complementary mouse resources that capture substantial genetic diversity and can demonstrate expansive phenotypic variation in heritable traits. These populations may be utilized to discover candidate genes and gene networks that moderate ethanol consumption and other ethanol-related traits.

**Methods:** We characterized ethanol consumption, preference and pharmacokinetics in the eight founder strains and ten CC strains in 12-hour drinking sessions during the dark phase of the circadian cycle.

**Results:** Ethanol consumption was found to be substantially heritable, both early in ethanol access and over a chronic intermittent access schedule. Ethanol pharmacokinetics were also found to be heritable; however, no association between strain-level ethanol consumption and pharmacokinetics was detected. The PWK/PhJ strain was found to be the highest drinking strain, with consumption substantially exceeding C57BL/6J, a strain commonly used as a model of “high” or “binge” drinking. Notably, we found strong evidence that sex moderated genetic effects on voluntary ethanol drinking.

**Conclusions:** Collectively, this research may serve as a foundation for expanded genetic study of ethanol consumption in the CC/DO and related populations; moreover, we have identified reference strains with extreme consumption phenotypes that effectively represent polygenic models of hazardous ethanol use.

## Introduction

Alcohol use disorder (AUD) is a psychiatric condition with a complex etiology and pathophysiology (Gilpin & Koob, 2008). Progress in understanding the causes of AUD and resulting biological dysfunction is required for discovery of pharmacotherapeutic and other interventions that may improve treatment outcomes. Genetics are a substantial etiological factor influencing ethanol use and AUD (Agrawal & Lynskey, 2008; Enoch & Goldman, 2001; Heath et al., 1991, 1997; Prescott et al., 1999; Reed et al., 1994; Swan et al., 1990; Verhulst et al., 2015a), with influences on traits that moderate onset of drinking and both quantity and patterns of drinking that may themselves predict risk of developing AUD (Chassin et al., 2002; Dash et al., 2020; Geels et al., 2012; Kalmus, 1967; McGue et al., 2001). While ethanol consumption and AUD may possess partially distinct genetic architecture (Sanchez-Roige et al., 2020), observed genetic correlations between these traits suggest that the identification of genetic factors affecting consumption may reveal mechanisms regulating AUD risk (Kranzler et al., 2019). However, the genes/gene networks implicated in traits that influence drinking patterns and the variants that drive their variation to potentially affect AUD vulnerability are largely undiscovered (Verhulst et al., 2015b). Consequently, major components of the biology that influences ethanol consumption and AUD risk remain poorly understood. Forward genetic, genome-wide approaches have the potential to implicate any variants/genes that influence ethanol consumption and ultimately contribute to a comprehensive understanding of AUD genetics.

The use of mouse populations for forward genetic screens allows for investigation of a broad range of quantitative traits under rigorous laboratory conditions. Some inbred strains of mice and some individual outbred mice will voluntarily consume ethanol at levels that produce relevant pharmacological effects including reward and intoxication; inter-strain and inter-individual variation, respectively, in this phenotype, is at least partially attributable to genetic variation (Belknap & Atkins, 2001; Grahame et al., 1999; Lopez et al., 2017; Rodriguez et al., 1994; Saba et al., 2011). Furthermore, several additional ethanol-related traits, including ethanol pharmacokinetic (PK) phenotypes, are readily measurable in mouse populations and are under the influence of genetic variation (Grisel et al., 2002). Reference inbred strain panels and outbred populations can be utilized for discovery of genes and gene networks that moderate these ethanol-related traits. The Collaborative Cross (CC), Diversity Outbred (DO), and Heterogenous Stock Collaborative Cross (HS-CC) mouse populations were founded from an advanced intercross of 8 distantly related strains (Chesler, 2014; Churchill et al., 2004; Iancu et al., 2010; Svenson et al., 2012). These populations capture a substantial amount of mouse genetic diversity. This key characteristic can expand the phenotypic range of the trait under study and may allow for expression of high levels of ethanol consumption (Chesler, 2014). This expanded variation may improve discovery of genes that drive ethanol-related behaviors and identify reference inbred strains with extreme phenotypes that may be utilized as improved models of AUD. To date, the HS-CC population has been successfully utilized as a selection base for selected breeding of alcohol preference for analysis of differential gene expression (Colville et al., 2017, 2018). Additionally, the CC/DO populations collectively may be utilized for forward genetic approaches that allow for quantitative trait locus (QTL) mapping of ethanol-related phenotypes in the DO and discovery of genetic correlations in the CC, including genetic correlation among transcript expression and ethanol-related phenotypes under study (Churchill et al., 2004; Svenson et al., 2012). The combination of these approaches may allow for nomination of candidate genes and gene networks that influence AUD related traits.

Here, we characterize ethanol consumption, preference and pharmacokinetics in both sexes of the eight founder strains of the CC/DO populations and ten CC strains. This approach will reveal the degree of heritable variation in ethanol consumption and metabolism, including potential interactions of genetic variation with sex. These data may serve as a foundation for expanded genetic study of ethanol-related traits in the CC/DO populations and reveal strains that are well suited to model heightened risk of developing an AUD-like state.

## Methods

### Subjects

Adult mice (ages detailed below) from the founder strains of the CC/DO populations and ten CC strains (see axis of figure 1 for strains, all mice obtained from The Jackson Laboratory, Bar Harbor, ME) were housed in a temperature- and humidity-controlled vivarium on a 12-hr light–dark cycle. Both sexes were tested in all strains. All mice were maintained on *ad libitum* mouse chow and water and housed in polycarbonate cages (30 × 8 cm) with wood-chip bedding (SANI-CHIPS, Montville, NJ), a paper nestlet and a red polycarbonate hut at a density of 1-4 mice per cage. All procedures were approved by the Binghamton University Institutional Animal Care and Use Committee and conducted in accordance with the National Institute of Health Guide for Care and Use of Laboratory Animals (National Research Council (US) Committee for the Update of the Guide for the Care and Use of Laboratory Animals, 2011).

**Figure 1.**
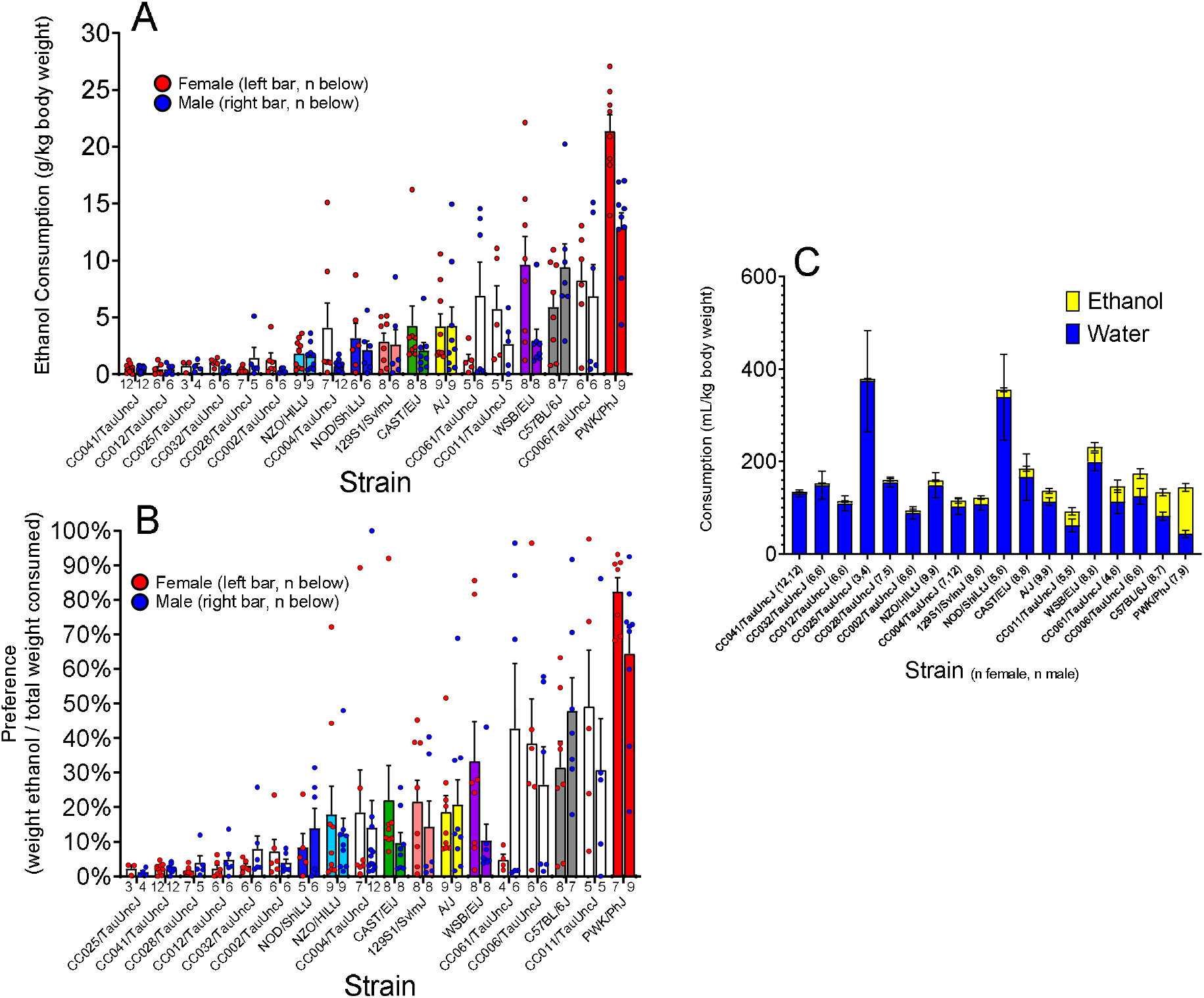
Ethanol consumption/preference and water consumption in founder and CC strains on day 2/3 of access. A) Ethanol consumption varied by strain with a heritability of H^2^ = 0.63, and sex interacted with strain. B) Preference of ethanol relative to concurrently available water varied by strain with a heritability of H^2^ = 0.49. C) Water consumption varied by strain and was negatively correlated with ethanol consumption.

### Ethanol Drinking

Ethanol drinking occurred in lickometer boxes (Scurry Mouse Dual Lickometer, Lafayette Instrument Company, Lafayette IN) housed in sound-attenuating isolation chambers. Test sessions began one hour into the dark phase (12/12 light cycle) and lasted for 12 hours. Two drinking bottles were provided, one with 20% (v/v) ethanol (190 proof diluted in water, Decon Labs Inc., PA USA) and the other with water (*ad lib* chow was also provided). Bottle start positions were counterbalanced between groups and alternated every session. Licks on both bottles were tracked for the whole session. The bottles were weighed immediately before and after the session to determine the amount consumed. Consumption of ethanol and water is expressed per kilogram of body weight.

### Ethanol Drinking Experiment 1: Three Consecutive Days

Mice of the founder strains and ten CC strains (see figure 1 for strains, 18 to 30 weeks of age) were tested for drinking over 3, consecutive 12-hour sessions. These mice were previously tested in a battery of behavioral testing that included: open field, chocolate-flavored Boost consumption in the same Scurry lickometer boxes, and operant tests of reversal learning or delay discounting. Mice were under food restriction during reversal learning/delay discounting. Upon completion, they were acclimated on *ad lib* diet for at least two weeks prior to ethanol drinking.

### Ethanol Drinking Experiment 2: Chronic Intermittent Access

A separate group of founder strain mice (10 weeks of age) were subject to a chronic intermittent access (CIA) schedule that included 4 days of consecutive 12-hour sessions in week 1 followed by a one day on, one day off schedule for three 12-hour sessions per week over 4 weeks. These mice were naïve to all testing prior to CIA drinking.

### Blood Ethanol Concentration

A separate group of founder strain mice (18 to 30 weeks of age), naïve to any form of ethanol exposure, were tested for blood ethanol concentration after intraperitoneal injection of 1.5 g/kg ethanol (20 mL/kg dose volume, ethanol diluted in saline to a concentration of 9.5% v/v). Blood was collected 30 minutes post-injection in all founder strains and (additionally) at 60 or 90 minutes post-injection for C57BL/6J and PWK/PhJ strains (different mice for each time point). Blood samples were spun to plasma and frozen at -80 C until assayed. BEC was determined by testing thawed samples using an Analox AM1 Alcohol Analyser (Analox Instruments, Amblecote, Stourbridge DY8 4HD, United Kingdom). The device was validated by ethanol standards at initial calibration and after every 5 samples.

### Data Analysis

Strain and sex were assessed by ANOVA. Heritability was estimated by partial eta-squared for strain. Genetic correlations were determined by performing Pearson’s correlations of strain means between each pair of traits. A p-value of less than 0.05 was considered significant. Where appropriate, post hoc analysis was performed by pairwise comparisons of estimated marginal means with Bonferroni p-value correction for multiple comparisons. Analysis was performed on SPSS version 27.

## Results

### Experiment 1: Ethanol Consumption and Preference

Ethanol consumption was assessed in mice with concurrent access to 20% ethanol and water during three, daily 12-hour sessions (drinking experiment 1). A mixed ANOVA revealed that ethanol consumption (g ethanol/kg body weight) differed by strain [F(17, 217)=25.8, p<0.001] and sex [F(1, 217)=5.7,p=0.017]. Consumption varied over session, an effect that was moderated by strain [F(34, 434)=2.7, p<0.001]. Subject-level correlational analyses revealed that consumption in sessions 2 and 3 were more similar to one another than either was to intake measured in session 1 (sessions 1 to 2: r=0.75; sessions 1 to 3: 0.71; sessions 2 to 3: r=0.82). For this reason, sessions 2 and 3 were averaged at the individual level, and this parameter was utilized in heritability analyses. Average intake during the latter 2 sessions exhibited main effects of strain [F(17, 221)=22.6, p<0.001], sex [F(1, 221)=5.4, p=0.021] and a strain-sex interaction [F(17, 221)=3.1, p<0.001]. The heritability of consumption was H^2^ =0.63 with strains demonstrating a broad range of consumption from near zero to approximately 17 g/kg/session in the PWK/PhJ strain (see figure 1A). Post hoc analyses indicated that the PWK/PhJ (p<0.001), WSB/EiJ (p<0.001), CC004/TauUncJ (p=0.039) and CC061/TauUncJ (p=0.007) strains demonstrated a sex effect.

Ethanol preference (volume of ethanol solution consumed as a percent of total liquid volume consumed) was analyzed using a mixed ANOVA, revealing an effect of strain [F(17, 214)=12.9, p<0.001] and a strain-session interaction [F(34, 428)=1.7, p=0.009]. Analysis of sessions 2 and 3 collapsed revealed an effect of strain with a heritability of H^2^ = 0.49 (see figure 1B).

Water consumption (adjusted for body weight) was assessed separately. Water consumption differed by strain [F(17, 214)=6.1, p<0.001], and strain interacted with session [F(34, 428)=1.5, p=0.03] (see figure 1 C). Water and ethanol intake, at the individual mouse level, were negatively correlated (r = -.30, p<0.001), suggesting higher ethanol intake associates with a *decrease* in water consumption.

### Strain Patterns of Drinking within Session

Lickometry was utilized to assess temporal patterns of ethanol intake within session 3. Average intake per hour was calculated and subjected to mixed ANOVA. A strain-hour-sex interaction (f[187, 2420)=1.2, p<0.034), a strain-hour interaction [F(187, 2420)=1.5, p<0.001], and a main effect of hour [F(11, 2420)= 3.6, p<0.05] were found, revealing that intake varies over the period of a session and this variability is influenced by strain and sex (see figure 2A). When analyzed within sex, strain was found to interact with hour only in males (f[187,1221]=1.8, p<0.001), suggesting the effect of strain on variability in drinking within session is more pronounced or specific to males. A main effect of strain [F(17, 220)=18.4, p<0.001], sex [F(1, 220)=5.1, p=0.025] and a strain-sex interaction [F(17, 220)=2.8, p<0.001] were also found, with patterns of strain and sex consumption commensurate with day 2/3 drinking as previously discussed (see figure 1A).

**Figure 2.**
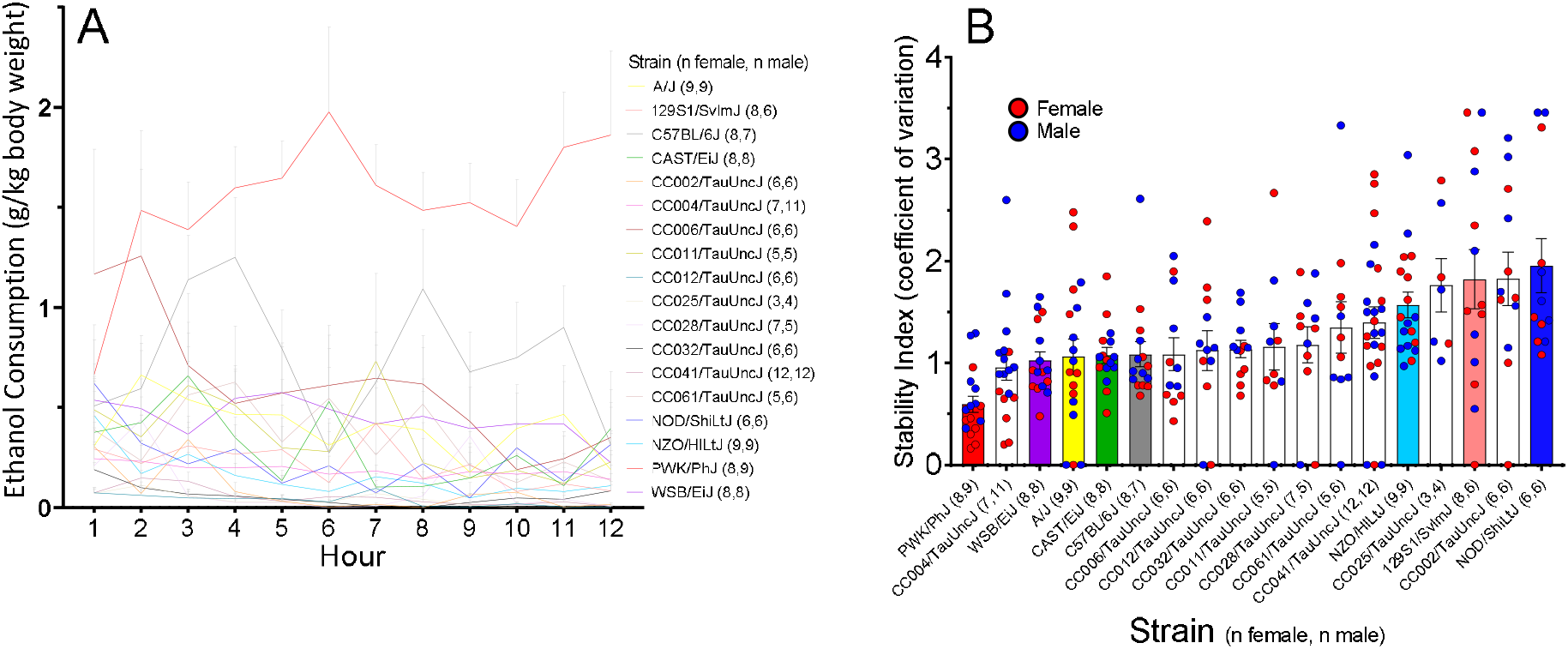
Temporal patterns of ethanol consumption within session on day 3. A) Ethanol consumption varied by hour in a strain-dependent manner. B) An index of drinking stability (coefficient of variation) revealed that strains varied in stability.

In order to produce an index of consumption variability within session, we calculated the coefficient of variation in the hour-by-hour ethanol drinking per mouse. This parameter differed by strain [F(17, 220)= 4.3, p<0.001], with PWK/PhJ mice demonstrating the lowest score and thus most temporally stable pattern of drinking (see figure 2B). This contrasts with the NOD/ShiLtJ strain, which demonstrates the lowest stability and low overall consumption. PWK/PhJ mice also contrast with the C57BL/6J strain, which ranks 2nd in consumption, yet has lower stability in drinking with an index approximately twice that of PWK/PhJ. C57Bl/6J mice drink in epochs separated by periods of substantially reduced consumption, as visualized in figure 2A.

### Experiment 2: Intermittent Ethanol Access

Consumption of and preference for ethanol (20%) was next studied over 5 weeks, with an intermittent access schedule in weeks 2 through 5; data was collapsed by week and subject to a mixed ANOVA. A main effect of strain [F(7, 14)=14.8, p<0.05] was found, indicating again that consumption differs as a function of strain. Additionally, a main effect of week [F(4, 56)= 3.3, p<0.05] indicates that consumption varied over time, with escalation being the most prevalent response (see figure 3A). Ethanol intake across the entire testing period exhibited a high heritability of 0.88, a figure substantially higher than the heritability estimate for intake measured only during the first week of consumption (0.64). Thus, genetic influences on intake seemed to grow as the period of access progressed.

**Figure 3.**
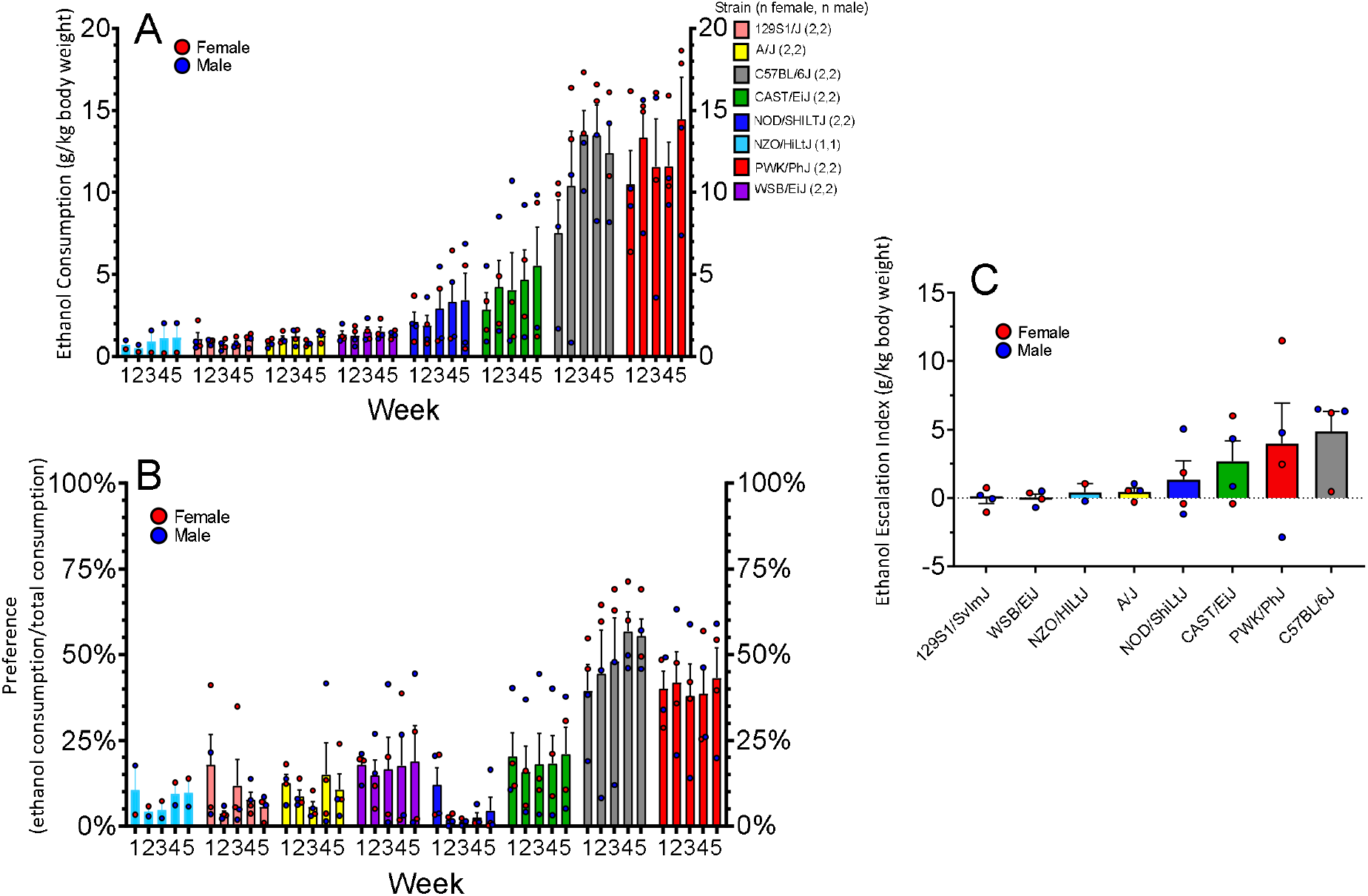
Ethanol consumption/preference and escalation of ethanol consumption in founder strains over 5 weeks of access, with a CIA schedule in weeks 2-5. A) Ethanol consumption varied by strain with a heritability of H^2^ = 0.88. B) Preference of ethanol relative to concurrently available water varied by strain with a heritability of H^2^ = 0.82. C) Escalation of intake varied by strain with a heritability of H^2^ = 0.41. Strains with high initial intake demonstrated subsequent escalation.

Analysis of ethanol preference also revealed main effects of strain [F(7, 14)=9.2, p<0.05] and week [F(4, 56)=2.8, p<0.05], with a heritability of 0.82 (see figure 3B).

To assess heritable variation in change in intake across the period of testing, an escalation index was calculated by subtracting the average intake in the last week of drinking from the first week at an individual mouse level. This phenotype exhibited a heritability estimate of H^2^ = 0.41 (see figure 3C). To assess the genetic relationship between escalation of drinking with initial intake, strain means for escalation were analyzed for correlation to mean consumption in the founder strains from experiment 1 (a different cohort of mice). The traits demonstrate a significant correlation [r = 0.93, p<0.05], indicating that strain-level consumption early in ethanol access was highly predictive of escalation over chronic intermittent access.

In order to assess the replicability of founder strain effects on ethanol drinking, strain means of session 2/3 from drinking experiment 1 and from drinking experiment 2 (different cohort of mice) were calculated and subject to Pearson’s correlation analyses. Ethanol drinking was found to be significantly correlated [r= 0.87, p<0.05], indicating that heritable variation in drinking was largely replicable between these two studies.

### Blood Ethanol Concentration

Blood ethanol concentrations (BEC) in trunk blood were determined in otherwise ethanol naïve mice from the founder strains, 30 minutes after an intraperitoneal ethanol injection. ANOVA revealed a main effect of strain [F(7, 30)=10.7, p<0.05] and sex [F(1, 30)=1.4, p<0.05], indicating that BEC demonstrates heritable variation, with a heritability estimate of H^2^ = 0.71 (see figure 4A). Genetic correlations between BEC at 30 min post-injection and ethanol drinking were assessed by determining the correlation between BEC strain means and consumption strain means. A significant correlation was not detected (r = -0.26, p>0.05) indicating that heritable variation in BEC was not predictive of ethanol drinking in this cohort.

**Figure 4.**
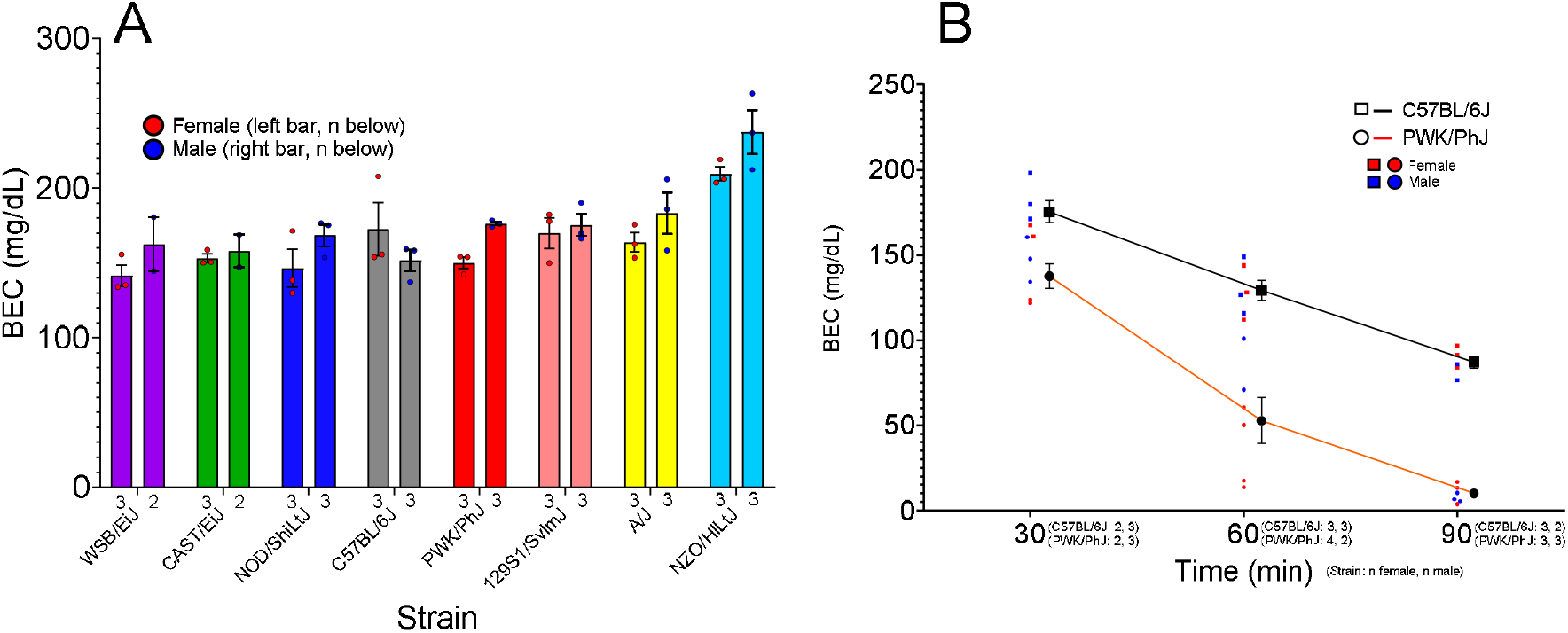
Blood ethanol concentration after ethanol injection in founder strains. A) Blood ethanol concentration varied by strain with a heritability of H^2^ = 0.71. B) Multiple post-injection timepoint measurements revealed that the PWK/PhJ strain clears ethanol at a greater rate compared to C57BL/6J.

The two highest drinking strains (C57Bl/6J and PWK/PhJ) were selected for further characterization of BEC at additional time points (60 or 90 min post-injection). An effect of time point was found, with BEC predictably declining over time. An effect of strain was found, with C57BL/6J demonstrating overall higher BEC at all time points (see figure 4B).

## Discussion

CC, DO and their founder strains are genetically diverse populations that may facilitate discovery of genes that influence ethanol consumption and preference, other ethanol-responsive behaviors and potential genetic relationships between these traits. We have characterized voluntary ethanol drinking and relevant ethanol pharmacokinetic phenotypes in the 8 founder strains and 10 CC strains to assess each for heritability and coinheritance of these traits. We find ethanol drinking to be substantially heritable, with a broad range of consumption between strains, ranging from near zero to a substantial 17 g/kg body weight (1.4 g/kg/hour) for the PWK/PhJ strain average with the female average exceeding 20 g/kg (1.7 g/kg/hour). Though procedural differences between published studies and this one make it challenging to make direct comparisons, the heritability and range of strain trait variation in the population reported here exceeds that of other inbred strain panels, e.g., BXD and AXB/BXA mice (Belknap & Atkins, 2001; Gill et al., 1996; Gill et al., 1998; Phillips et al., 1994; Rodriguez et al., 1995), and outbred populations (Belknap & Atkins, 2001; Phillips et al., 1998). These results suggest that the CC/ DO populations are well suited for forward genetics studies of voluntary ethanol consumption and include reference strains that are polygenic models of extreme ethanol consumption phenotypes, generating new opportunities for the study of the mechanisms driving hazardous drinking.

Ethanol drinking was studied in two experiments; the first characterized drinking in founder strains and ten CC strains during three consecutive nightly sessions and the second utilized founder strains tested under CIA conditions, for a total of 16 sessions spanning 5 weeks. With respect to the former, ethanol drinking exhibited a highly stable pattern of substantial heritability by the 2^nd^ and 3^rd^ consumption test. Moreover, when comparing intake in the two experiments, strain-level variation in intake was found to be replicable, despite that the cohorts of mice in these two experiments were tested at different ages and with different experimental histories. This suggests that ethanol consumption traits are robust to differences in age and experimental history. Analysis of preference, intake of ethanol solution relative to concurrently available water, revealed similar results although heritability was somewhat lower for this trait. Given that heritable variation in ethanol consumption can be reliably measured in just a few sessions, this phenotype could be amenable to high throughput testing of large populations of DO mice, enabling subsequent genome-wide or transcriptome-wide analyses to identify molecular variants that could underlie behavioral differences.

Despite the longer period of time required for testing, CIA ethanol drinking is useful because one can observe escalation of ethanol intake over consecutive drinking epochs (Hwa et al., 2011; Melendez, 2011; Rosenwasser et al., 2013; Spear, 2020). Consequently, escalation of drinking during CIA may reveal additional and unique genetic information not captured when only examining short-term access to ethanol. We did observe strain-dependent escalation of ethanol intake. However, strain-level intake in the 3-day access test was highly predictive of the degree of subsequent escalation. CIA was only tested in the founder strains, and it is possible that further characterization of CC or DO mice may reveal heritable variation in escalation that is independent of intake early in access. Notably, the heritability of drinking under the CIA schedule was substantially greater relative to that observed in the short-term access test. This suggests that extended access under the CIA schedule may improve the precision of phenotypic measurement and, in turn, the discovery of QTL and genetic correlations. Overall, the present results indicate that these populations are suitable for chronic ethanol drinking procedures. Inter-strain relationships remain stable over multi-week timescales while the higher drinking strains escalate intake, suggesting that this procedure is suited to investigate the genetics of chronic and high-level ethanol drinking.

We examined temporal patterns of drinking within-session by utilizing lickometry data. This analysis may reveal relevant information not captured when analysis is done at the session level or collapsed across multiple sessions. Critically, ethanol pharmacokinetics and resulting pharmacodynamics may be greatly influenced by the temporal patterns of drinking, in addition to total intake. These patterns may range from punctuated intake, with periods of drinking followed by periods of reduced intake, to more stable intake over the session. We observed heritable variation in patterns of intake. The PWK/PhJ strain demonstrated the most stable pattern of intake, in addition to the being the highest drinking strain, while others – like C57BL/6J – exhibited substantial within-session variability in the amount of intake. These results suggest that genetically influenced patterns of ethanol intake may have important consequences for ethanol PK. Peak plasma concentration and time at concentration are important variables that may influence ethanol pharmacodynamics, including influence on brain physiology and behavior. The present data suggest that variation in drinking patterns may cause differences in peak, and time under, plasma concentration and this measure may be utilized to complement total intake variables in genetic and neurobiological studies.

Physiological factors that influence the pharmacokinetic profile of the blood ethanol concentration are likely under genetic influence in mice, as they are in humans. Genetic influences on ethanol metabolism are risk factors for AUD and may directly influence levels of ethanol drinking (Eckey et al., 1990; Frank et al., 2012; Gelernter et al., 2018; Jorgenson et al., 2017; Otto et al., 2013; Park et al., 2013; Quillen et al., 2014; Takeuchi et al., 2011; Walters et al., 2018). We found that BEC after non-contingent injection was heritable in the founder strains. The range of observed BEC at the strain-level was not substantial, with most strains falling between 150 and 175 mg/dL, with the exception of NZO/HlLtJ which exceeded 200. We did not find evidence to suggest that strain differences in ethanol PK influenced strain-level ethanol drinking (strain-level correlation r = -0.27, p = 0.52). These results suggest that heritable pharmacodynamic factors may be influential in determining differences in founder strain drinking.

Sex differences in ethanol consumption and other ethanol responsive traits are important etiological factors for AUD (Agabio et al., 2017). The present results indicate that sex interacts with genetic background to influence ethanol consumption. Consumption of ethanol appears approximately equivalent between sexes in many strains whereas some demonstrate large sex differences. The PWK/PhJ strain, in particular, demonstrates a robust sex effect with females drinking approximately 60% more than males on average. Sex differences in BEC after ethanol injection were also discovered, with males tending to achieve higher BEC than females. Sex differences have been previously reported in mice; however, the direction of effect varies by study and may be influenced by route of administration (Baraona et al., 2001; Desroches et al., 1995; Frezza et al., 1990; Middaugh et al., 1992). Collectively, these results suggest that the CC/DO population may be utilized to discover genes that interact with biological sex to influence ethanol consumption and PK. This information may prove critical in elucidating the biological factors that drive sex differences in ethanol consumption.

In addition to discovery of candidate genes and genetic correlations, characterization of ethanol consumption in the founder and CC populations may reveal reference strains with extreme phenotypes that can be utilized as improved models of ethanol consumption. This is particularly likely with the founder and CC strains due to the large degree of genetic variability captured in these populations (Philip et al., 2011; Threadgill & Churchill, 2012). Previously, the HS-CC population has been successfully utilized as a base for selective breeding for alcohol preference, with multi-generational breeding producing a population with high ethanol preference and consumption (Colville et al., 2017, 2018). The present study has demonstrated that extreme ethanol phenotypes are present in the founder panel and that founders, CC and DO populations collectively may capture extreme ethanol phenotypes without the need for additional, multi-generational selective breeding. Ethanol drinking studies that utilize mice are frequently based on the C57BL/6J strain. This strain is known to voluntarily consume ethanol at a quantity that can produce pharmacodynamic effects on the central nervous system and resulting behavioral changes (Crabbe, 2014). The present results indicate that the PWK/PhJ strain demonstrates an extreme ethanol consumption phenotype that reliably exceeds consumption by C57BL/6J. High ethanol consumption by PWK/PhJ may indicate that this strain is suited to model ethanol consumption traits that are informative for problematic drinking and AUD. Although no CC strains were found to substantially exceed PWK/PhJ or C57BL/6J (the highest drinking CC strain is approximately equivalent to C57BL/6J), the range of founder strain drinking suggests that CC strain panel and DO population may capture a broad range of ethanol consumption, with consumption exceeding PWK/PhJ possible, although this will only be revealed by further characterization of an expanded CC or DO sample.

Extensive characterization of ethanol PK in the C57BL/6J and PWK/PhJ strains revealed that PWK/PhJ may have metabolic or excretion phenotypes that support enhanced clearance of ethanol. This may influence strain differences in BEC levels after voluntary drinking. Although PWK/PhJ may clear ethanol at a faster rate, they drink substantially more. Whether they achieve greater BEC levels under voluntary drinking remains unknown and requires additional studies. Analysis of temporal patterns of drinking also revealed strain differences in the stability of drinking, with PWK/PhJ demonstrating a more stable pattern relative to C57BL/6J. These patterns may also have consequences for peak blood concentration and duration of blood levels. Additional studies are required to better elucidate the differences between PWK/PhJ and C57BL/6J however, these results suggest that PWK/PhJ may be useful to expand genetic and neurobiological research of high ethanol consumption and sex differences in this trait.

These data have established that ethanol consumption is a substantially heritable trait in CC and their founder strains, perhaps serving as a foundation for expanded characterization of ethanol consumption in CC strains and DO mice. The use of DO mice for high-resolution QTL mapping combined with characterization of CC strains for genetic correlation studies provides opportunities to identify genes and gene networks that moderate ethanol consumption. Additionally, these data have revealed that PWK/PhJ is a high drinking strain with a prominent sex difference that may be utilized to study both acute and chronic ethanol consumption. Collectively, this research indicates that the CC/DO populations and their founders are a valuable resource for investigating the genetics and neurobiology of ethanol consumption.

## Acknowledgements

These studies were supported, in part, by Public Health Service grants T32-AA025606 (JDJ and JRB), P50-DA039841 and P50-AA017823 (JDJ). We would like to thank Lauren Bailey and Barbara Force, for their assistance and technical support in this research.

## Notes

### Competing Interest Statement

The authors have declared no competing interest.

## References

Agabio, R., Pisanu, C., Gessa, G. L., & Franconi, F. (2017). Sex Differences in Alcohol Use Disorder. Current Medicinal Chemistry, 24(24). https://doi.org/10.2174/0929867323666161202092908

Agrawal, A., & Lynskey, M. T. (2008). Are there genetic influences on addiction: Evidence from family, adoption and twin studies. Addiction, 103(7), 1069–1081. https://doi.org/10.1111/j.1360-0443.2008.02213.x

Baraona, E., Abittan, C. S., Dohmen, K., Moretti, M., Pozzato, G., Chayes, Z. W., Schaefer, C., & Lieber, C. S. (2001). Gender Differences in Pharmacokinetics of Alcohol. Alcoholism: Clinical and Experimental Research, 25(4), 502–507. https://doi.org/10.1111/j.1530-0277.2001.tb02242.x

Belknap, J. K., & Atkins, A. L. (2001). The replicability of QTLs for murine alcohol preference drinking behavior across eight independent studies. Mammalian Genome: Official Journal of the International Mammalian Genome Society, 12(12), 893–899. https://doi.org/10.1007/s00335-001-2074-2

Chassin, L., Pitts, S. C., & Prost, J. (2002). Binge drinking trajectories from adolescence to emerging adulthood in a high-risk sample: Predictors and substance abuse outcomes. Journal of Consulting and Clinical Psychology, 70(1), 67–78. https://doi.org/10.1037/0022-006X.70.1.67

Chesler, E. J. (2014). Out of the bottleneck: The Diversity Outcross and Collaborative Cross mouse populations in behavioral genetics research. Mammalian Genome, 25(1–2), 3–11. https://doi.org/10.1007/s00335-013-9492-9

Churchill, G. A., Airey, D. C., Allayee, H., Angel, J. M., Attie, A. D., Beatty, J., Beavis, W. D., Belknap, J. K., Bennett, B., Berrettini, W., Bleich, A., Bogue, M., Broman, K. W., Buck, K. J., Buckler, E., Burmeister, M., Chesler, E. J., Cheverud, J. M., Clapcote, S., … Zou, F. (2004). The Collaborative Cross, a community resource for the genetic analysis of complex traits. Nature Genetics, 36(11), 1133–1137. https://doi.org/10.1038/ng1104-1133

Colville Alexandre M., Iancu, O. D., Lockwood, D. R., Darakjian, P., McWeeney, S. K., Searles, R., Zheng, C., & Hitzemann, R. (2018). Regional Differences and Similarities in the Brain Transcriptome for Mice Selected for Ethanol Preference From HS-CC Founders. Frontiers in Genetics, 9. https://doi.org/10.3389/fgene.2018.00300

Colville Alexandre M., Iancu, O. D., Oberbeck, D. L., Darakjian, P., Zheng, C. L., Walter, N. a. R., Harrington, C. A., Searles, R., McWeeney, S., & Hitzemann, R. (2017). Effects of selection for ethanol preference on gene expression in the nucleus accumbens of HS-CC mice. Genes, Brain, and Behavior, 16(4), 462–471. https://doi.org/10.1111/gbb.12367

Crabbe, J. C. (2014). Rodent Models of Genetic Contributions to Motivation to Abuse Alcohol. Nebraska Symposium on Motivation. Nebraska Symposium on Motivation, 61, 5–29.

Dash, G. F., Davis, C. N., Martin, N. G., Statham, D. J., Lynskey, M. T., & Slutske, W. S. (2020). High-Intensity Drinking in Adult Australian Twins. Alcoholism, Clinical and Experimental Research, 44(2), 522–531. https://doi.org/10.1111/acer.14262

Desroches, D., Orevillo, C., & Verina, D. (1995). Sex-and strain-related differences in first-pass alcohol metabolism in mice. Alcohol, 12(3), 221–226. https://doi.org/10.1016/0741-8329(94)00098-X

Eckey, R., Agarwal, D. P., & Goedde, H. W. (1990). [Genetically-induced variability of alcohol metabolism and its effect on drinking behavior and predisposition to alcoholism]. Zeitschrift Fur Rechtsmedizin. Journal of Legal Medicine, 103(3), 169–190. https://doi.org/10.1007/BF00207339

Enoch, M.-A., & Goldman, D. (2001). The genetics of alcoholism and alcohol abuse. Current Psychiatry Reports, 3(2), 144–151. https://doi.org/10.1007/s11920-001-0012-3

Frank, J., Cichon, S., Treutlein, J., Ridinger, M., Mattheisen, M., Hoffmann, P., Herms, S., Wodarz, N., Soyka, M., Zill, P., Maier, W., Mössner, R., Gaebel, W., Dahmen, N., Scherbaum, N., Schmäl, C., Steffens, M., Lucae, S., Ising, M., … Rietschel, M. (2012). Genome-wide significant association between alcohol dependence and a variant in the ADH gene cluster. Addiction Biology, 17(1), 171–180. https://doi.org/10.1111/j.1369-1600.2011.00395.x

Frezza, M., di Padova, C., Pozzato, G., Terpin, M., Baraona, E., & Lieber, C. S. (1990). High Blood Alcohol Levels in Women. New England Journal of Medicine, 322(2), 95–99. https://doi.org/10.1056/NEJM199001113220205

Geels, L. M., Bartels, M., van Beijsterveldt, T. C. E. M., Willemsen, G., van der Aa, N., Boomsma, D. I., & Vink, J. M. (2012). Trends in adolescent alcohol use: Effects of age, sex and cohort on prevalence and heritability. Addiction (Abingdon, England), 107(3), 518–527. https://doi.org/10.1111/j.1360-0443.2011.03612.x

Gelernter, J., Zhou, H., Nuñez, Y. Z., Mutirangura, A., Malison, R. T., & Kalayasiri, R. (2018). Genomewide Association Study of Alcohol Dependence and Related Traits in a Thai Population. Alcoholism: Clinical and Experimental Research, 42(5), 861–868. https://doi.org/10.1111/acer.13614

Gill, K., Liu, Y., & Deitrich, R. A. (1996). Voluntary Alcohol Consumption in BXD Recombinant Inbred Mice: Relationship to Alcohol Metabolism. Alcoholism: Clinical and Experimental Research, 20(1), 185–190. https://doi.org/10.1111/j.1530-0277.1996.tb01063.x

Gill, Kathryn, Desaulniers, N., Desjardins, P., & Lake, K. (1998). Alcohol preference in AXB/BXA recombinant inbred mice: Gender differences and gender-specific quantitative trait loci. Mammalian Genome, 9(12), 929–935. https://doi.org/10.1007/s003359900902

Gilpin, N. W., & Koob, G. F. (2008). Neurobiology of Alcohol Dependence. Alcohol Research & Health, 31(3), 185–195.

Grahame, N. J., Li, T. K., & Lumeng, L. (1999). Selective breeding for high and low alcohol preference in mice. Behavior Genetics, 29(1), 47–57. https://doi.org/10.1023/a:1021489922751

Grisel, J. E., Metten, P., Wenger, C. D., Merrill, C. M., & Crabbe, J. C. (2002). Mapping of Quantitative Trait Loci Underlying Ethanol Metabolism in BXD Recombinant Inbred Mouse Strains. Alcoholism: Clinical and Experimental Research, 26(5), 610–616. https://doi.org/10.1111/j.1530-0277.2002.tb02582.x

Heath, A. C., Bucholz, K. K., Madden, P. a. F., Dinwiddie, S. H., Slutske, W. S., Bierut, L. J., Statham, D. J., Dunne, M. P., Whitfield, J. B., & Martin, N. G. (1997). Genetic and environmental contributions to alcohol dependence risk in a national twin sample: Consistency of findings in women and men. Psychological Medicine, 27(6), 1381–1396. https://doi.org/10.1017/S0033291797005643

Heath, A. C., Meyer, J., Jardine, R., & Martin, N. G. (1991). The inheritance of alcohol consumption patterns in a general population twin sample: II. Determinants of consumption frequency and quantity consumed. Journal of Studies on Alcohol, 52(5), 425–433. https://doi.org/10.15288/jsa.1991.52.425

Hwa, L. S., Chu, A., Levinson, S. A., Kayyali, T. M., DeBold, J. F., & Miczek, K. A. (2011). Persistent Escalation of Alcohol Drinking in C57BL/6J Mice With Intermittent Access to 20% Ethanol. Alcoholism: Clinical and Experimental Research, 35(11), 1938–1947. https://doi.org/10.1111/j.1530-0277.2011.01545.x

Iancu, O. D., Darakjian, P., Walter, N. A., Malmanger, B., Oberbeck, D., Belknap, J., McWeeney, S., & Hitzemann, R. (2010). Genetic diversity and striatal gene networks: Focus on the heterogeneous stock-collaborative cross (HS-CC) mouse. BMC Genomics, 11(1), 585. https://doi.org/10.1186/1471-2164-11-585

Jorgenson, E., Thai, K. K., Hoffmann, T. J., Sakoda, L. C., Kvale, M. N., Banda, Y., Schaefer, C., Risch, N., Mertens, J., Weisner, C., & Choquet, H. (2017). Genetic contributors to variation in alcohol consumption vary by race/ethnicity in a large multi-ethnic genome-wide association study. Molecular Psychiatry, 22(9), 1359–1367. https://doi.org/10.1038/mp.2017.101

Kalmus, H. (1967). Inheritance of Drinking Behavior: A Study on Intelligence, Personality, and Use of Alcohol of Adult Twins. Journal of the Royal Statistical Society: Series A (General), 130(4), 589–589. https://doi.org/10.2307/2982543

Kranzler, H. R., Zhou, H., Kember, R. L., Vickers Smith, R., Justice, A. C., Damrauer, S., Tsao, P. S., Klarin, D., Baras, A., Reid, J., Overton, J., Rader, D. J., Cheng, Z., Tate, J. P., Becker, W. C., Concato, J., Xu, K., Polimanti, R., Zhao, H., & Gelernter, J. (2019). Genome-wide association study of alcohol consumption and use disorder in 274,424 individuals from multiple populations. Nature Communications, 10(1), 1499. https://doi.org/10.1038/s41467-019-09480-8

Lopez, M. F., Miles, M. F., Williams, R. W., & Becker, H. C. (2017). Variable effects of chronic intermittent ethanol exposure on ethanol drinking in a genetically diverse mouse cohort. Alcohol (Fayetteville, N.Y.), 58, 73–82. https://doi.org/10.1016/j.alcohol.2016.09.003

McGue, M., Iacono, W. G., Legrand, L. N., & Elkins, I. (2001). Origins and Consequences of Age at First Drink. II. Familial Risk and Heritability. Alcoholism: Clinical and Experimental Research, 25(8), 1166–1173. https://doi.org/10.1111/j.1530-0277.2001.tb02331.x

Melendez, R. I. (2011). Intermittent (every-other-day) drinking induces rapid escalation of ethanol intake and preference in adolescent and adult C57BL/6J mice. Alcoholism, Clinical and Experimental Research, 35(4), 652–658. https://doi.org/10.1111/j.1530-0277.2010.01383.x

Middaugh, L. D., Frackelton, W. F., Boggan, W. O., Onofrio, A., & Shepherd, C. L. (1992). Gender differences in the effects of ethanol on C57BL/6 mice. Alcohol, 9(3), 257–260. https://doi.org/10.1016/0741-8329(92)90062-F

Otto, J. M., Hendershot, C. S., Collins, S. E., Liang, T., & Wall, T. L. (2013). Association of the ALDH1A1*2 Promoter Polymorphism With Alcohol Phenotypes in Young Adults With or Without ALDH2*2. Alcoholism: Clinical and Experimental Research, 37(1), 164–169. https://doi.org/10.1111/j.1530-0277.2012.01835.x

Park, B. L., Kim, J. W., Cheong, H. S., Kim, L. H., Lee, B. C., Seo, C. H., Kang, T.-C., Nam, Y.-W., Kim, G.-B., Shin, H. D., & Choi, I.-G. (2013). Extended genetic effects of ADH cluster genes on the risk of alcohol dependence: From GWAS to replication. Human Genetics, 132(6), 657–668. https://doi.org/10.1007/s00439-013-1281-8

Philip, V. M., Sokoloff, G., Ackert-Bicknell, C. L., Striz, M., Branstetter, L., Beckmann, M. A., Spence, J. S., Jackson, B. L., Galloway, L. D., Barker, P., Wymore, A. M., Hunsicker, P. R., Durtschi, D. C., Shaw, G. S., Shinpock, S., Manly, K. F., Miller, D. R., Donohue, K. D., Culiat, C. T., … Chesler, E. J. (2011). Genetic analysis in the Collaborative Cross breeding population. Genome Research, 21(8), 1223– 1238. https://doi.org/10.1101/gr.113886.110

Phillips, T. J., Belknap, J. K., Buck, K. J., & Cunningham, C. L. (1998). Genes on mouse chromosomes 2 and 9 determine variation in ethanol consumption. Mammalian Genome: Official Journal of the International Mammalian Genome Society, 9(12), 936–941. https://doi.org/10.1007/s003359900903

Phillips, Tamara J., Crabbe, J. C., Metten, P., & Belknap, J. K. (1994). Localization of Genes Affecting Alcohol Drinking in Mice. Alcoholism: Clinical and Experimental Research, 18(4), 931–941. https://doi.org/10.1111/j.1530-0277.1994.tb00062.x

Prescott, C. A., Aggen, S. H., & Kendler, K. S. (1999). Sex Differences in the Sources of Genetic Liability to Alcohol Abuse and Dependence in a Population-Based Sample of U.S. Twins. Alcoholism: Clinical and Experimental Research, 23(7), 1136–1144. https://doi.org/10.1111/j.1530-0277.1999.tb04270.x

Quillen, E. E., Chen, X.-D., Almasy, L., Yang, F., He, H., Li, X., Wang, X.-Y., Liu, T.-Q., Hao, W., Deng, H.-W., Kranzler, H. R., & Gelernter, J. (2014). ALDH2 is associated to alcohol dependence and is the major genetic determinant of “daily maximum drinks” in a GWAS study of an isolated rural chinese sample. American Journal of Medical Genetics Part B: Neuropsychiatric Genetics, 165(2), 103–110. https://doi.org/10.1002/ajmg.b.32213

Reed, T., Slemenda, C. W., Viken, R. J., Christian, J. C., Carmelli, D., & Fabsitz, R. R. (1994). Correlations of alcohol consumption with related covariates and heritability estimates in older adult males over a 14-to 18-year period: The NHLBI Twin Study. Alcoholism, Clinical and Experimental Research, 18(3), 702–710. https://doi.org/10.1111/j.1530-0277.1994.tb00934.x

Rodriguez, L. A., Plomin, R., Blizard, D. A., Jones, B. C., & McClearn, G. E. (1994). Alcohol acceptance, preference, and sensitivity in mice. I. Quantitative genetic analysis using BXD recombinant inbred strains. Alcoholism, Clinical and Experimental Research, 18(6), 1416–1422. https://doi.org/10.1111/j.1530-0277.1994.tb01444.x

Rodriguez, Lawrence A., Plomin, R., Blizard, D. A., Jones, B. C., & McClearn, G. E. (1995). Alcohol Acceptance, Preference, and Sensitivity in Mice. II. Quantitative Trait Loci Mapping Analysis Using BXD Recombinant Inbred Strains. Alcoholism: Clinical and Experimental Research, 19(2), 367–373. https://doi.org/10.1111/j.1530-0277.1995.tb01517.x

Rosenwasser, A. M., Fixaris, M. C., Crabbe, J. C., Brooks, P. C., & Ascheid, S. (2013). Escalation of intake under intermittent ethanol access in diverse mouse genotypes. Addiction Biology, 18(3), 496– 507. https://doi.org/10.1111/j.1369-1600.2012.00481.x

Saba, L. M., Bennett, B., Hoffman, P. L., Barcomb, K., Ishii, T., Kechris, K., & Tabakoff, B. (2011). A systems genetic analysis of alcohol drinking by mice, rats and men: Influence of brain GABAergic transmission. Neuropharmacology, 60(7–8), 1269–1280. https://doi.org/10.1016/j.neuropharm.2010.12.019

Sanchez-Roige, S., Palmer, A. A., & Clarke, T.-K. (2020). Recent Efforts to Dissect the Genetic Basis of Alcohol Use and Abuse. Biological Psychiatry, 87(7), 609–618. https://doi.org/10.1016/j.biopsych.2019.09.011

Spear, L. P. (2020). Timing Eclipses Amount: The Critical Importance of Intermittency in Alcohol Exposure Effects. Alcoholism: Clinical and Experimental Research, 44(4), 806–813. https://doi.org/10.1111/acer.14307

Svenson, K. L., Gatti, D. M., Valdar, W., Welsh, C. E., Cheng, R., Chesler, E. J., Palmer, A. A., McMillan, L., & Churchill, G. A. (2012). High-Resolution Genetic Mapping Using the Mouse Diversity Outbred Population. Genetics, 190(2), 437–447. https://doi.org/10.1534/genetics.111.132597

Swan, G. E., Carmelli, D., Rosenman, R. H., Fabsitz, R. R., & Christian, J. C. (1990). Smoking and alcohol consumption in adult male twins: Genetic heritability and shared environmental influences. Journal of Substance Abuse, 2(1), 39–50. https://doi.org/10.1016/s0899-3289(05)80044-6

Takeuchi, F., Isono, M., Nabika, T., Katsuya, T., Sugiyama, T., Yamaguchi, S., Kobayashi, S., Ogihara, T., Yamori, Y., Fujioka, A., & Kato, N. (2011). Confirmation of ALDH2 as a Major Locus of Drinking Behavior and of Its Variants Regulating Multiple Metabolic Phenotypes in a Japanese Population. Circulation Journal, 75(4), 911–918. https://doi.org/10.1253/circj.CJ-10-0774

Threadgill, D. W., & Churchill, G. A. (2012). Ten Years of the Collaborative Cross. Genetics, 190(2), 291– 294. https://doi.org/10.1534/genetics.111.138032

Verhulst, B., Neale, M. C., & Kendler, K. S. (2015a). The heritability of alcohol use disorders: A meta-analysis of twin and adoption studies. Psychological Medicine, 45(5), 1061–1072. https://doi.org/10.1017/S0033291714002165

Verhulst, B., Neale, M. C., & Kendler, K. S. (2015b). The heritability of alcohol use disorders: A meta-analysis of twin and adoption studies. Psychological Medicine, 45(5), 1061–1072. https://doi.org/10.1017/S0033291714002165

Walters, R. K., Polimanti, R., Johnson, E. C., McClintick, J. N., Adams, M. J., Adkins, A. E., Aliev, F., Bacanu, S.-A., Batzler, A., Bertelsen, S., Biernacka, J. M., Bigdeli, T. B., Chen, L.-S., Clarke, T.-K., Chou, Y.-L., Degenhardt, F., Docherty, A. R., Edwards, A. C., Fontanillas, P., … Agrawal, A. (2018). Trans-ancestral GWAS of alcohol dependence reveals common genetic underpinnings with psychiatric disorders. Nature Neuroscience, 21(12), 1656–1669. https://doi.org/10.1038/s41593-018-0275-1

